# Multi-view confounder detection for biomedical studies

**DOI:** 10.1101/2022.10.14.512210

**Authors:** Markus Wallner, Nicolas Kersten, Nico Pfeifer

## Abstract

In many biomedical studies an important first step is checking for confounding factors. For association studies, confounding can for example be caused by ethnic differences in the case and control groups. In many other settings there might be confounding factors like batch effects or founder effects that also need to be detected and controlled for^1^. Detecting confounding for data from one data source is well established (e.g., genomics data). Since more and more studies are now based on data from multiple data modalities (e.g., multi-omics), we evaluated whether multi-view confounder detection can benefit from state-of-the-art methods for multi-view data integration. Especially for clustering of multi-omics data, it has been shown that these methods can perform better than methods that treat the data modalities separately^2^. Our results show that multi-view confounder analysis is possible and that building on multi-view data integration methods is better than treating the different data modalities separately.

---

When analyzing the genetic basis for common and complex diseases, confounding factors have to be detected and controlled for to decrease the likelihood of finding false positive associations^3^. If one also has additional data modalities like methylation data, then these data have to be controlled for confounding, too^4^. If the signal for confounding is weak and distributed among several data modalities (e.g., genetic data and methylation data), methods for joint detection of confounding might be more sensitive compared to methods that check for confounding separately. Similar advantages have been shown for cancer subtype detection based on multi-omics data^2^ as well as a synthetic benchmark data set^5^. To the best of our knowledge, multi-view confounder detection has never been evaluated systematically. Therefore, we devised a similar synthetic benchmark data set to the one introduced for multi-omics clustering by Rappoport and Shamir (2019)^5^ and evaluated the performance of different ways to detect confounding. In addition to simulated data sets, we also performed similar analyses on real cancer data.

In order to compare the different approaches, we devised a setting in which all methods can be used in a similar way to detect the confounding. The main hypothesis is that if samples cluster into two groups that are not independent of the potential confounder, then this confounder could influence any analysis of the data and should therefore be accounted for. Therefore, we tested whether there was a significant correlation between the resulting cluster labels of each method and the labels of the potential confounding factor by performing a permutation test based on chi-squared tests. The only difference in the evaluated methods is how to devise the clusters or how to handle the results from the tests based on the clusters.

As shown in a benchmark by Rappoport and Shamir in 2018^2^, there are some key differences in performance between available multi-view clustering algorithms. In order to evaluate the ability to detect multi-view confounding factors, we developed multi-view confounding tests (Multi-Contest) based on several state-of-the-art methods like rMKL-LPP^6^ (Multi-Contest^rMKL-LPP^), NEMO^5^ (Multi-Contest^NEMO^), COCA^7^ (Multi-Contest^COCA^), as well as two simple logical operations (Multi-Contest^and-P^, Multi-Contest^or-P^). The clustering by COCA as well as the logical operations are based on single omics K-means clustering. For Multi-Contest^or-P^ one and for Multi-Contest^and-P^ all omics have to show a significant p-value to detect the potential confounding factor in question. Based on these detections and the ground truth we calculated Matthews correlation coefficient (MCC) to compare the performance in the different simulation settings.

The first simulation setting contains two cluster centers. The simulated samples are represented by points normally distributed around the respective cluster centers. The omics are then generated by adding normal distributed noise (Gaussian noise) to these simulated sample points. We increased the spread of samples around the cluster center as well as the spread of omics around their sample point to test how the performance is influenced. For all these settings we tested different numbers of omics and different strengths of the confounding. Fig. 1 summarizes the performance of the five tested methods for the first simulation setting (sensitivity and specificity plots in Supplementary Fig. 1). Overall, a higher spread of omics/samples leads to lower MCC values. The spread of samples around the cluster centers seems to have a stronger impact than the spread of omics around the samples. For the four different settings of sample and omics spread in this setup, Multi-Contest^rMKL-LPP^ and Multi-Contest^NEMO^ outperform the other methods. The two logical operations perform worst, with Multi-Contest^and-P^ showing significantly lower MCC values.

**Figure 1.**
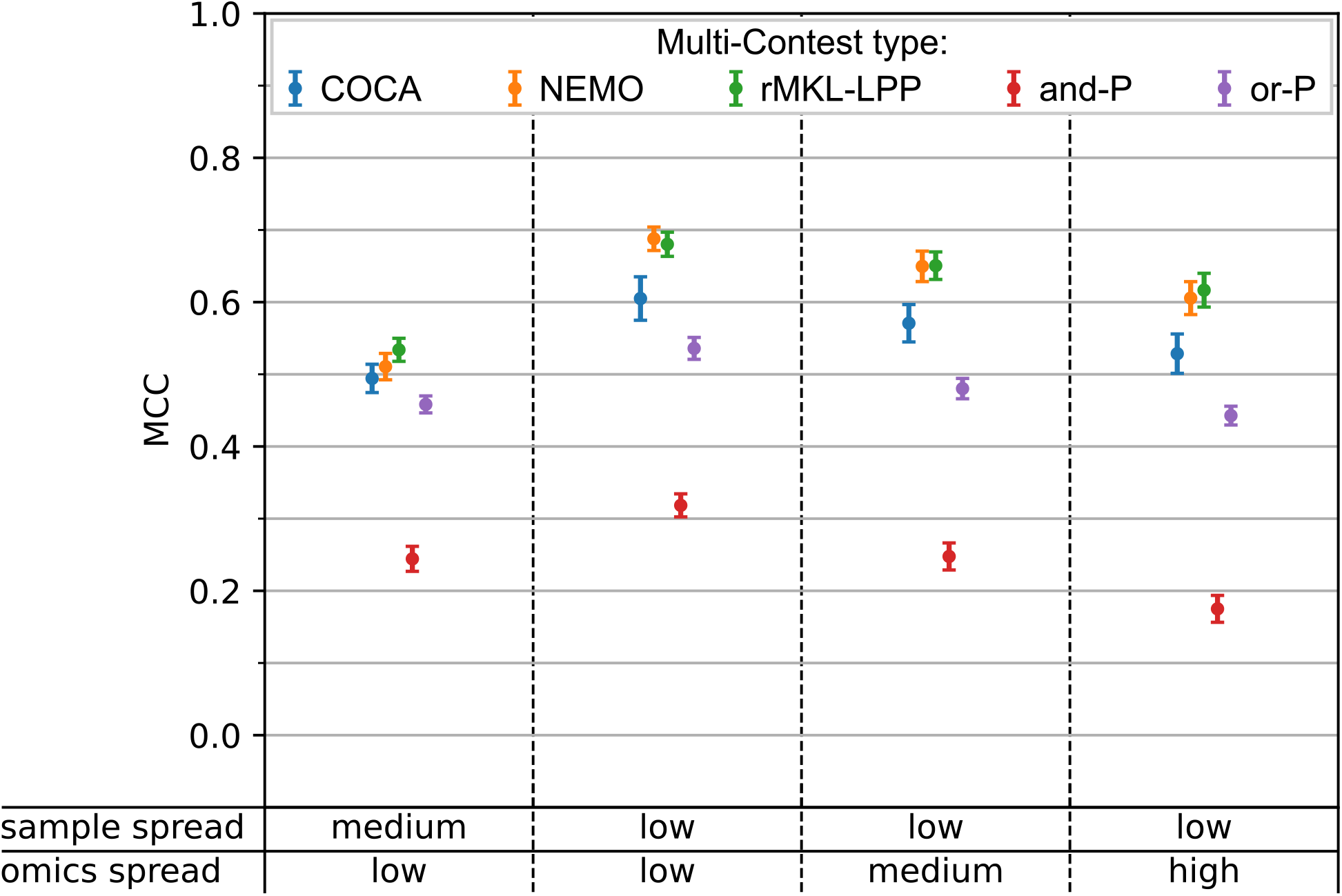
MCC values for the simulation settings with two cluster centers. Results are shown as means *±* SEM of Matthews Correlation Coefficient (MCC) values from different number of omics and strength of simulated confounding (triplicates for each combination, total n = 36). Different combinations of sample spread and omics spread were used to test how the methods perform on a less clear cluster structure.

To evaluate the different methods in a more complex setup, we extended the simulation setting by introducing a third cluster in which all clusters have the same distance to each other. The results for this setting are shown in Fig. 2 (sensitivity and specificity plots in Supplementary Fig. 2). Here Multi-Contest^rMKL-LPP^ and Multi-Contest^NEMO^ show the highest MCC values as well. The distance to the next best method (Multi-Contest^COCA^) is larger than in the previous simulation setting. Multi-Contest^or-P^ shows notably worse performance in this setting, while Multi-Contest^NEMO^ and Multi-Contest^rMKL-LPP^ achieve similar MCC values compared to the first setting.

**Figure 2.**
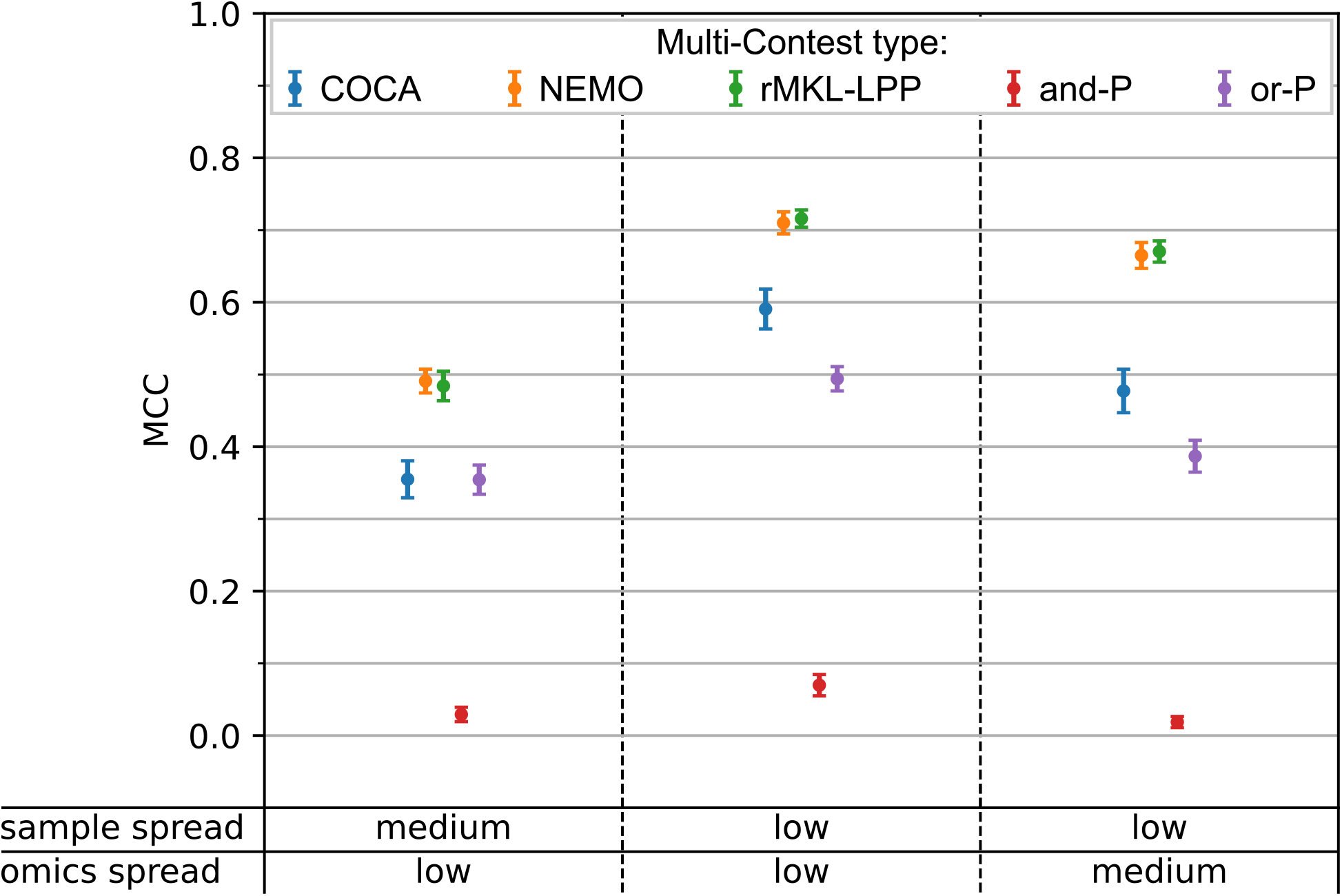
MCC values for the simulation settings with three cluster centers. Results are shown as means *±* SEM of Matthews Correlation Coefficient (MCC) values from different strength of simulated confounding with ten omics (triplicates for each strength, total n = 24). Different combinations of sample spread and omics spread were used to test how the methods perform on less a less clear cluster structure.

Up to this point all simulated omics of a data set were based on the same underlying structure without omics that were not affected by the confounding factor. In order to create a more refined simulation, we changed the simulation setting such that only a portion of the omics is influenced by a confounding signal with a similar strength as before (now called signal omics / SO). For the remaining omics we reduced the distance between cluster centers and increased the spread for samples around these centers as well as the spread of omics around the sample points. These omics are now called noise omics (NO). The total number of omics per sample was fixed to ten. We tested this setting with 2, 3, and 5 SO and two different cluster distances for the NO each. The resulting MCC values are summarized in Fig. 3 (sensitivity and specificity plots in Supplementary Fig. 3). In this setting the performance of Multi-Contest^NEMO^ (MCC) seems to depend strongly on the number of signal omics. The other methods (except for Multi-Contest^and-P^) show a trend of higher MCC values with more signal omics as well, but the order of the methods does not change. Only Multi-Contest^NEMO^ improves from second worst with a high amount of noise omics to best/shared best MCC values with an equal amount of signal omics and noise omics. Considering all variations of this setting Multi-Contest^rMKL-LPP^ shows the most robust performance.

**Figure 3.**
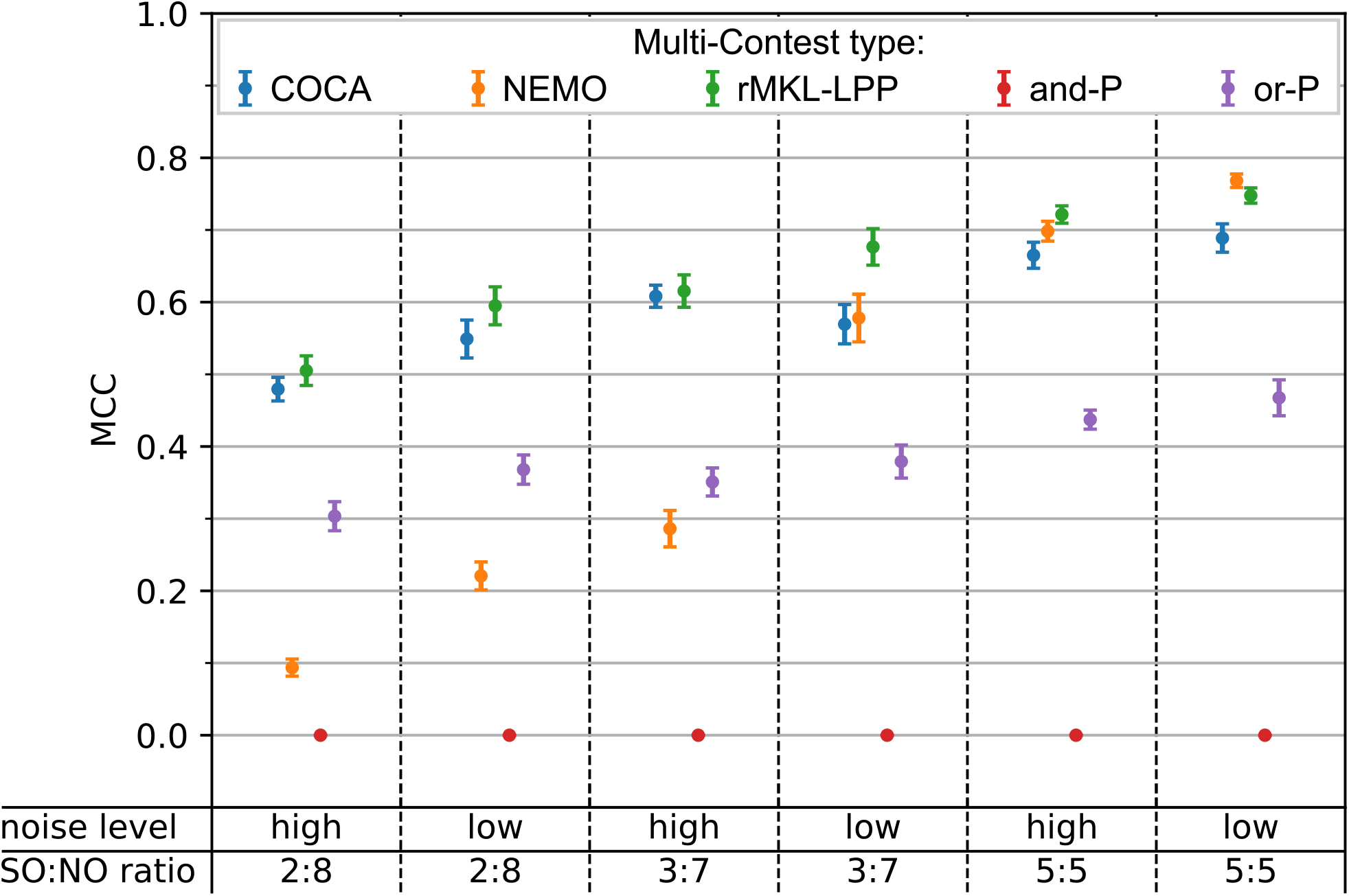
MCC values for simulated data sets containing signal omics and noise omics. Results are shown as means *±* SEM of Matthews Correlation Coefficient (MCC) values from different strength of simulated confounding with ten omics (triplicates for each strength, total n = 12). The signal omics (SO) to noise omics (NO) ratio (SO:NO ratio) as well as the noise level used for the simulation are indicated at the bottom of the figure.

Finally we evaluated the methods on three TCGA data sets with the following cancer types: Glioblastoma Multiforme (GBM), Liver Hepatocellular Carcinoma (LIHC), and Lung Squamous Cell Carcinoma (LUSC). For each data set gene expression, methylation, and miRNA expression data was included. Additionally we used the gender information as potential confounding factor and compared confounding tests for individual omics clusterings to the previously described Multi-Contest settings. The resulting p-values for Multi-Contest^or-P^ (where a significant test result in one of the omics is sufficient) were adjusted for multiple testing using Bonferroni correction. All resulting p-values are summarized in Table 1.

**Table 1.** Resulting p-values from testing gender labels as potential confounding factor for the three cancer data sets Liver Hepatocellular Carcinoma (LIHC), Glioblastoma Multiforme (GBM), Lung Squamous Cell Carcinoma (LUSC), and a reduced GBM data set (GBM-red.) obtained from GBM by removing the expression data. The expression, methylation, and miRNA results are based on single omics K-means clusterings. For Multi-Contest^and-P^ the highest p-value of the individual omics is reported (because all have to give a significant result). For Multi-Contest^or-P^ the lowest p-value of the single omics is reported (with adjustment for multiple testing) because a significant result in one of the tested omics is sufficient. Multi-Contest^COCA^ combines the information of these three single omics clusterings, while Multi-Contest^NEMO^ and Multi-Contest^rMKL-LPP^ do not rely on those single omics clusterings. Values below the signifcance threshold of *α* = 0.05 are shown in bold face.

| Method | p-value |  |  |  |
| --- | --- | --- | --- | --- |
|  | LIHC | GBM | GBM-red. | LUSC |
| expression | **$< 10^{-3}$** | **0.013** | - | 0.073 |
| methylation | **0.031** | 0.517 | 0.517 | 0.261 |
| miRNA | **0.041** | 1.000 | 1.000 | 0.727 |
| Multi-Contest<sup>and-P</sup> | **0.041** | 1.000 | 1.000 | 0.727 |
| Multi-Contest<sup>or-P</sup> | **$< 3 \cdot 10^{-3}$** | **0.039** | 1.000 | 0.219 |
| Multi-Contest<sup>COCA</sup> | **$< 10^{-3}$** | 0.104 | 1.000 | 0.073 |
| Multi-Contest<sup>NEMO</sup> | **$< 10^{-3}$** | **$< 10^{-3}$** | **$< 10^{-3}$** | 0.060 |
| Multi-Contest<sup>rMKL-LPP</sup> | **0.040** | **0.010** | **0.042** | 0.097 |

For the LIHC data set the clustering of the baseline methods as well as Multi-Contest^COCA^, Multi-Contest^NEMO^, and Multi-Contest^rMKL-LPP^ lead to a positive test result. In this data set the confounding effect is strong enough to be even detectable based on the individual omics data. For this cancer type previously a male to female ratio of 2.5 was observed indicating some kind of gender bias^8^. With ongoing research, a number of molecular mechanisms that are leading to the observed sex-bias have been discovered^9^–12.

For the GBM data set only the test of confounding based on gene expression, Multi-Contest^or-P^, Multi-Contest^NEMO^, and Multi-Contest^rMKL-LPP^ resulted in a significant p-value and therefore indicated gender as a confounding factor. Since significant confounding could in this case already be detected by only analyzing gene expression data, an additional evaluation was performed on the remaining two data types that did not show significant confounding in the single-omics setting (GBM-red.). In the GBM-red. data set the more sophisticated methods Multi-Contest^NEMO^ and Multi-Contest^rMKL-LPP^ still lead to a significant test result. Previous studies showed that for some GBM subtypes a sex-bias is present^13^. A recent study also reported sex-specific molecular subtypes of glioblastoma multiforme^14^. This matches with the positive test result based on gene expression, Multi-Contest^or-P^, Multi-Contest^NEMO^, and Multi-Contest^rMKL-LPP^. In the GBM-red. example the confounding was still detectable based on Multi-Contest^NEMO^ and Multi-Contest^rMKL-LPP^ even without the expression data. This is an example for a data set where the representation from the integration of multiple omics using consensus clustering (COCA) does not indicate the presence of confounding, while the more sophisticated representations based on NEMO and rMKL-LPP enable a positive test result.

Lung squamous cell carcinoma is a type of non-small cell lung carcinoma (NSCLC). For NSCLC male gender was confirmed to be an independent unfavorable prognostic indicator for survival^15^. Other studies also reported the influence of gender on targeted treatments^16^,17 as well as gender-specific somatic alterations that influence prognostic biomarkers^10^. The sex-bias that is reported in the literature was not observed with any of the tested methods for this data set, while some of the advanced methods showed a trend towards significance. Since a ground truth is not available here, it is not possible to decide if the confounding was not detected or if it was not present in this specific data set.

All results from the simulated data sets show, that the confounder detection based on the more sophisticated multi-omics clustering methods NEMO and rMKL-LPP outperforms the approaches based on the individual omics. For the TCGA data sets we did observe that the confounding in the GBM-red. data set was still detectable by using NEMO and rMKL-LPP representations even though the omic that was most influenced by the confounding factor was removed. This was not possible using the other tested methods. This indicates that when the confounding effect is less pronounced, then methods that consider the omics data jointly are better suited to detect the confounding than methods which evaluate the different data modalities separately.

## Methods

### Data simulation

In order to evaluate the performance of different approaches to reveal the presence of confounding factors, different data sets were simulated. With simulated data sets the ground truth about cluster affiliation of samples and the presence of confounding is known. The basis for the simulation settings is a simulation used by Rappoport and Shamir to test the performance of multi-omics clustering methods (including NEMO)^5^.

All simulated data sets contain 300 samples and for each setting 300 data sets were simulated. Each sample gets labeled with a gender label (‘m’ or ‘f’) as confounding factor, 150 samples per label. With no confounding present in these data sets no difference in gender label proportions in the clusters would be expected. To introduce confounding, the probability for samples with a specific gender label to be assigned to a certain cluster was changed in the simulation. All cluster centers (CC1 - CC5 are shown in Table 2. The most basic simulations contain two cluster centers in R^10^ at CC1 and CC2. The samples were then randomly assigned to the two cluster centers. For the samples labeled with ‘f’ both clusters had a probability of 0.5, for the ones labeled with ‘m’ the probability for the first cluster center was increased to different levels in the range 0.55 - 0.70 (step size 0.05) to simulate different levels of confounding. Each sample gets assigned a sample point based on a multivariate normal distribution with the identity matrix *I* as covariance matrix around the cluster centers. Individual omics profiles are now generated by adding normal noise *𝒩* (*µ* = 0, Σ = 2*I*) to the sample points. Data sets containing 2, 5, and 10 omics were created to model different sizes of data sets.

**Table 2.** Cluster centers (CC) used for simulation of synthetic data sets.

| Cluster center | Coordinates in $\mathbb{R}^{10}$ |
| --- | --- |
| CC1 | (2, 2, 2, 2, 2, 0, 0, 0, 0, 0) |
| CC2 | (0, 0, 0, 0, 0, 0, 0, 0, 0, 0) |
| CC3 | (1, 1, 1, 1, 1, 1.732, 1.732, 1.732, 1.732, 1.732) |
| CC4 | (1, 1, 1, 1, 1, 0, 0, 0, 0, 0) |
| CC5 | (0.5, 0.5, 0.5, 0.5, 0.5, 0, 0, 0, 0, 0) |

**Table 3.** Covariance matrices used in simulation (sample/omics spread) of synthetic data sets based on labels used in result graphs.

| Label in graph | sample spread | omics spread |
| --- | --- | --- |
| low | 1I | 2I |
| medium | 2I | 3I |
| high | - | 4I |

To test the influence of the spread of samples around the cluster centers and omics around sample points we changed the covariance matrices used for omics generation to *3I* and *4I* and for the final setting the covariance matrix for the sample point generation to *2I* while keeping the omics generation with *2I*.

In the next step the simulation setting was extended to three cluster centers by adding a third cluster center (CC3). In the process the probabilities for the cluster assignment were adjusted as well. The samples labeled with ‘f’ have equal probability for all three clusters. For the samples labeled with ‘m’ the probability to get assigned to the first cluster was set in the range 0.15 - 0.5 (step size 0.05) while the remaining probability was split evenly between the remaining two clusters.

With this setting only data sets containing ten omics were simulated. For the spread of samples around the cluster centers or omics around the sample points, three settings were tested. First the default covariance matrices as in the first setting was used, secondly *2I* for the samples around the cluster centers, and lastly *3I* was used for the noise generating the omics from the sample points.

Up to this point, all omics of one data set were based on the same underlying structure (position of the sample points around the cluster centers). To better simulate different information content and structure in different omics a third setting was created. Here data sets with two cluster centers and 10 omics were created with a number of omics (2, 3, 5) containing a rather clear cluster separation (signal omics / SO) and the remaining omics where the cluster centers are closer together and the spread is increased (noise omics / NO). With this general approach two noise levels were used. The signal omics were created with the same settings as used in the first two cluster setting. For the first noise level (low) the cluster centers are CC4 and CC2 with covariance matrix for sample points equal to *2I* and for omics generation equal to *3I*.

For the second noise level (high) the cluster centers CC5 and CC2 were used with the same covariance matrices.

### Real omics preprocessing

In order to test the performance of the different methods on real omics data we used the Glioblastoma Multiforme (GBM), Liver Hepatocellular Carcinoma (LIHC), Lung Squamous Cell Carcinoma (LUSC) preprocessed data sets that were previously used to benchmark multi-omics clustering algorithms^2^ and are based on TCGA data sets (downloaded from http://acgt.cs.tau.ac.il/multi_omic_benchmark/download.html). On these data sets the method-specific preprocessings were applied as described in the benchmark^2^. Each data set contains gene expression, DNA methylation and miRNA expression data. All samples that were not labeled as primary tumor were dropped. For each data set all patients that are not present in all three omics were dropped (resulting number of samples: GBM:274, LIHC:369, LUSC:341).

For K-means clustering, rMKL-LPP and Nemo, the data was filtered to exclude features with zero variance and all features based on sequencing data were log-transformed. The data was further reduced for K-means clustering such that only the 2000 highest variance gene expression and methylation features were selected. Finally all selected features were normalized to have a mean of zero and a standard deviation of 1. In addition to the complete real omics data sets, we created a reduced version of the GBM data (GBM-red.) by removing the gene expression data modality.

For these preprocessing steps we used the R functions from the NEMO GitHub page (https://github.com/Shamir-Lab/NEMO).

### K-means

K-means clusterings were performed on the single omics data sets. Here the K-means function of the scikit-learn Python package (sklearn.cluster.KMeans) was used. The best value for the number of clusters k was selected by average silhouette score for a predefined range of k (2 <= k <= 15) (sklearn.metrics.silhouette_score).

### Cluster-Of-Cluster Analysis (COCA)

In order to integrate the clusterings based on the single omics types we used Cluster Of Clusters Analysis (COCA). This method is widely used in cancer studies^7^,18,19 since its first introduction^20^. The goal of this method is to combine the information of the single omics clusterings into one consensus clustering. The R package “coca” was used to perform the consensus clustering step based on a matrix of clusters created form the single omics k-means clusterings. The maximum number of iterations (B) was set to 2000. All other parameters were used with their default values.

### Neighborhood-based multi-omics clustering (NEMO)

NEMO is a simple similarity-based multi-omics clustering approach^2^. The clustering is based on inter-patient similarity matrices for the different omics that get integrated into one matrix before the final clustering step. We used the R package as described in the publication^2^. All parameters were kept at default values.

### rMKL-LPP

Regularized Multiple Kernel Learning with Locality Preserving Projections (rMKL-LPP) is an unsupervised joint dimensionality reduction method that was first introduced in 2015^6^. rMKL-LPP performs a non-linear dimensionality reduction by using a kernelized representation of the sample-similarities for each data type. By learning a weighted linear combination of these kernel matrices, multi-omics data is integrated and projected to the n-dimensional space. Sample clusters are subsequently computed by applying the K-means algorithm to the dimensionality-reduced projection of the samples. To determine the number of clusters k, the average silhouette score metric is used for a predefined range of k (2 <= k <= 15).

The input for rMKL-LPP are similarity matrices, here computed using the radial basis function (RBF) kernel.

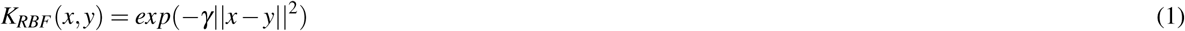

For all data sets (simulated and real cancer data) the parameter *γ* of the RBF kernel was set to the general rule of thumb 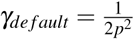, with *p* being the number of features^21^.

For the rMKL calculation, two parameters were selected, *k*_*N*_ the number of neighbors and *n* the number of dimensions to which the samples will be projected. For *k*_*N*_ the default value of nine was used for all computations. For the simulated data sets the number of dimensions *n* was set to ten. For the cancer data sets, the number of dimensions was set to five (the default value for web-rMKL^22^).

### Test for presence of confounding factors

The test for the presence of confounding is based on cluster labels as well as labels for potential confounding factors. We used the chi-squared test for independence with cluster assignments and potential confounder labels (‘f’/’m’ in the simulated data sets). Previous studies reported the possibility of inaccurate approximation of the statistic and a resulting overestimation of significance^2^,23. Because of this we decided to perform permutation tests as they were suggested previously^2^,5.

For the permutation test we randomly permuted the cluster labels of the samples. We performed 1*e*3 permutations at a time until the 95% confidence interval did not cross 0.05 or the maximum number of 1*e*5 iterations was reached. The confidence intervals were computed based on the number of permutations resulting in a greater or equal chi-squared statistic compared to the original labeling. For this the binomtest of scipy (scipy.stats.binomtest) was used. For the chi-squared test we used a function from python scipy (scipy.stats.chi2_contingency). If the resulting p-value is significant with an alpha cutoff of 0.05 the tested factor is counted as a confounding factor.

For the most basic evaluation we used the K-means clusterings of the single omics and required either one (or-P) or all omics (and-P) to give a significant result to count the tested factor as confounder. This results in multiple tests per data set and therefore bonferroni multiple testing correction was applied (using statsmodels.stats.multitest.multipletests) for the or-P setting. For the more advanced methods (COCA, NEMO, rMKL-LPP) the result is only one clustering so no multiple testing correction was required. We called the tests based on these clusterings Multi-Contest^COCA^, Multi-Contest^NEMO^, and Multi-Contest^rMKL-LPP^. For the simulated data sets the ground truth for the presence of confounding is calculated based on the known true cluster assignments used in the simulation.

### Matthews correlation coefficient

To compare the confounder detection performance of the different methods over the different simulation settings we calculated Matthews correlation coefficient (MCC) (calculated with sklearn.metrics.matthews_corrcoef). The MCC values are calculated for each data set and then combined into plots. One plot for the simple settings containing two clusters, one for the settings with three clusters, and one for the settings with different strength of confounding in the omics. For the simple two cluster settings the different covariance matrix combinations (sample/omics spread) are plotted as separate points while the different number of omics and the different strength of confounding is combined into mean and standard error. For the settings with three clusters an equivalent plot is created (here no combination of different numbers of omics in the data set were simulated). For the final simulation setting the different combinations of covariance matrices and weakness level get plotted as separate points.

## Acknowledgements

The results shown here are in whole or part based upon data generated by the TCGA Research Network: https://www.cancer.gov/tcga. M.W. and N.P. acknowledge funding from ZIV Consortium (Center for Innovative Care, digital@bw, 42-04HV.MED(18)/25/1). N.K. acknowledges funding from the Ministery of Economic Affairs, Labour and Housing of Baden-Württemberg (WM BW) within the ‘PRIMO’ project (reference number 3-4332.62-HSG/84).

## Author contributions statement

Conceptualization, N.P. and M.W.; data simulation and processing, M.W.; implementation and statistical analysis, M.W. and N.K.; project supervision, N.P. and N.K.; All authors contributed equally to result interpretation, manuscript preparation and manuscript revision.

## Additional information

The authors declare no conflicts of interest.

## Notes

### Competing Interest Statement

The authors have declared no competing interest.

## References

1. Carlson, J. M. et al. Impact of pre-adapted HIV transmission. Nat. Medicine 22, 606–613, DOI: http://doi.org/10.1038/nm.4100 (2016).

2. Rappoport, N. & Shamir, R. Multi-omic and multi-view clustering algorithms: review and cancer benchmark. Nucleic acids research 46, 10546–10562, DOI: http://doi.org/10.1093/nar/gky889 (2018). Publisher: Oxford University Press.

3. Pourhoseingholi, M. A., Baghestani, A. R. & Vahedi, M. How to control confounding effects by statistical analysis. Gastroenterol Hepatol Bed Bench 5, 79–83 (2012). Type: Journal Article.

4. Barfield, R. T. et al. Accounting for population stratification in DNA methylation studies. Genet. Epidemiol 38, 231–41, DOI: http://doi.org/10.1002/gepi.21789 (2014). xType: Journal Article.

5. Rappoport, N. & Shamir, R. NEMO: Cancer subtyping by integration of partial multi-omic data. Bioinformatics 35, 3348–3356, DOI: http://doi.org/10.1093/bioinformatics/btz058 (2019). Publisher: Oxford University Press.

6. Speicher, N. K. & Pfeifer, N. Integrating different data types by regularized unsupervised multiple kernel learning with application to cancer subtype discovery. Bioinforma. (Oxford, England) 31, i268–i275, DOI: http://doi.org/10.1093/bioinformatics/btv244 (2015).

7. Hoadley, K. A. et al. Multiplatform analysis of 12 cancer types reveals molecular classification within and across tissues of origin. Cell 158, 929–944, DOI: http://doi.org/10.1016/j.cell.2014.06.049 (2014). Type: Journal Article.

8. Keng, V. W. et al. Sex bias occurrence of hepatocellular carcinoma in Poly7 molecular subclass is associated with EGFR. Hepatology 57, 120–130, DOI: http://doi.org/10.1002/hep.26004 (2013). Publisher: Wiley Online Library.

9. Li, Z., Tuteja, G., Schug, J. & Kaestner, K. H. Foxa1 and Foxa2 are essential for sexual dimorphism in liver cancer. Cell 148, 72–83, DOI: http://doi.org/10.1016/j.cell.2011.11.026 (2012). Publisher: Elsevier.

10. Li, C. H., Haider, S., Shiah, Y.-J., Thai, K. & Boutros, P. C. Sex differences in cancer driver genes and biomarkers. Cancer research 78, 5527–5537, DOI: http://doi.org/10.1158/0008-5472.CAN-18-0362 (2018). Publisher: AACR.

11. Li, C. H. et al. Sex differences in oncogenic mutational processes. Nat. Commun. 11, 4330, DOI: http://doi.org/10.1038/s41467-020-17359-2 (2020).

12. Warrington, N. M. et al. The cyclic AMP pathway is a sex-specific modifier of glioma risk in type I neurofibromatosis patients. Cancer research 75, 16–21, DOI: http://doi.org/10.1158/0008-5472.CAN-14-1891 (2015). Publisher: AACR.

13. Sun, T., Plutynski, A., Ward, S. & Rubin, J. B. An integrative view on sex differences in brain tumors. Cell. molecular life sciences 72, 3323–3342, DOI: http://doi.org/10.1007/s00018-015-1930-2 (2015). Publisher: Springer.

14. Yang, W. et al. Sex differences in GBM revealed by analysis of patient imaging, transcriptome, and survival data. Sci. translational medicine 11, DOI: http://doi.org/10.1126/scitranslmed.aao5253 (2019). Publisher: American Association for the Advancement of Science.

15. Visbal, A. L. et al. Gender differences in non–small-cell lung cancer survival: an analysis of 4,618 patients diagnosed between 1997 and 2002. The Annals thoracic surgery 78, 209–215, DOI: http://doi.org/10.1016/j.athoracsur.2003.11.021 (2004). Publisher: Elsevier.

16. Pinto, J. A. et al. Gender and outcomes in non-small cell lung cancer: an old prognostic variable comes back for targeted therapy and immunotherapy? ESMO open 3, e000344, DOI: http://doi.org/10.1136/esmoopen-2018-000344 (2018). Publisher: Elsevier.

17. Schmetzer, O. & Flörcken, A. Sex differences in the drug therapy for oncologic diseases. Sex gender differences pharmacology 411–442, DOI: http://doi.org/10.1007/978-3-642-30726-3_19 (2013). Publisher: Springer.

18. Aure, M. R. et al. Integrative clustering reveals a novel split in the luminal A subtype of breast cancer with impact on outcome. Breast Cancer Res. 19, 1–18, DOI: http://doi.org/10.1186/s13058-017-0812-y (2017). Publisher: Springer.

19. Cabassi, A. & Kirk, P. D. Multiple kernel learning for integrative consensus clustering of omic datasets. Bioinformatics 36, 4789–4796, DOI: http://doi.org/10.1093/bioinformatics/btaa593 (2020). Publisher: Oxford University Press.

20. Network, T. C. G. A. Comprehensive molecular portraits of human breast tumours. Nature 490, 61 (2012). Publisher: Nature Publishing Group.

21. Gärtner, T., Flach, P. A., Kowalczyk, A. & Smola, A. J. Multi-instance kernels. In ICML, 7 (2002). Issue: 3.

22. Röder, B., Kersten, N., Herr, M., Speicher, N. K. & Pfeifer, N. web-rMKL: a web server for dimensionality reduction and sample clustering of multi-view data based on unsupervised multiple kernel learning. Nucleic Acids Res. 47, W605–W609, DOI: http://doi.org/10.1093/nar/gkz422 (2019).

23. Vandin, F., Papoutsaki, A., Raphael, B. J. & Upfal, E. Accurate computation of survival statistics in genome-wide studies. PLoS computational biology 11, e1004071, DOI: http://doi.org/10.1371/journal.pcbi.1004071 (2015). Publisher: Public Library of Science San Francisco, CA USA.

