## Supplementary Figures for "Multi-view confounder detection for biomedical studies"

Supplementary Materials

### Supplementary Figures

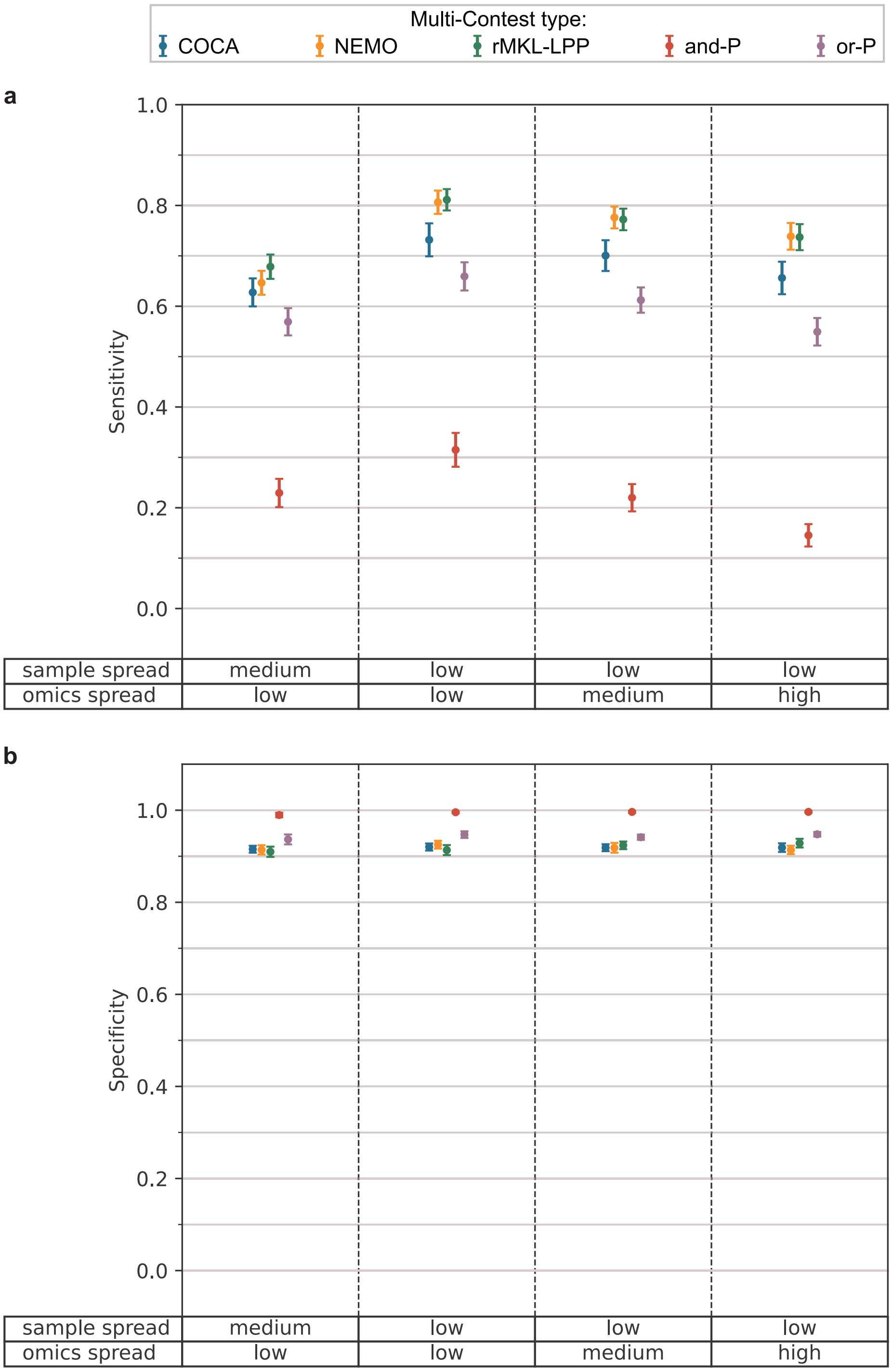


**Supplementary Figure 1:** Performance measures of Multi-Contest in simulated data sets with two cluster centers. The sensitivity (**a**) and specificity (**b**) are shown as means ± SEM from different number of omics and strength of simulated confounding (triplicates for each combination, total n = 36). Different combinations of sample spread and omics spread were used to test how the methods perform on less a less clear cluster structure.

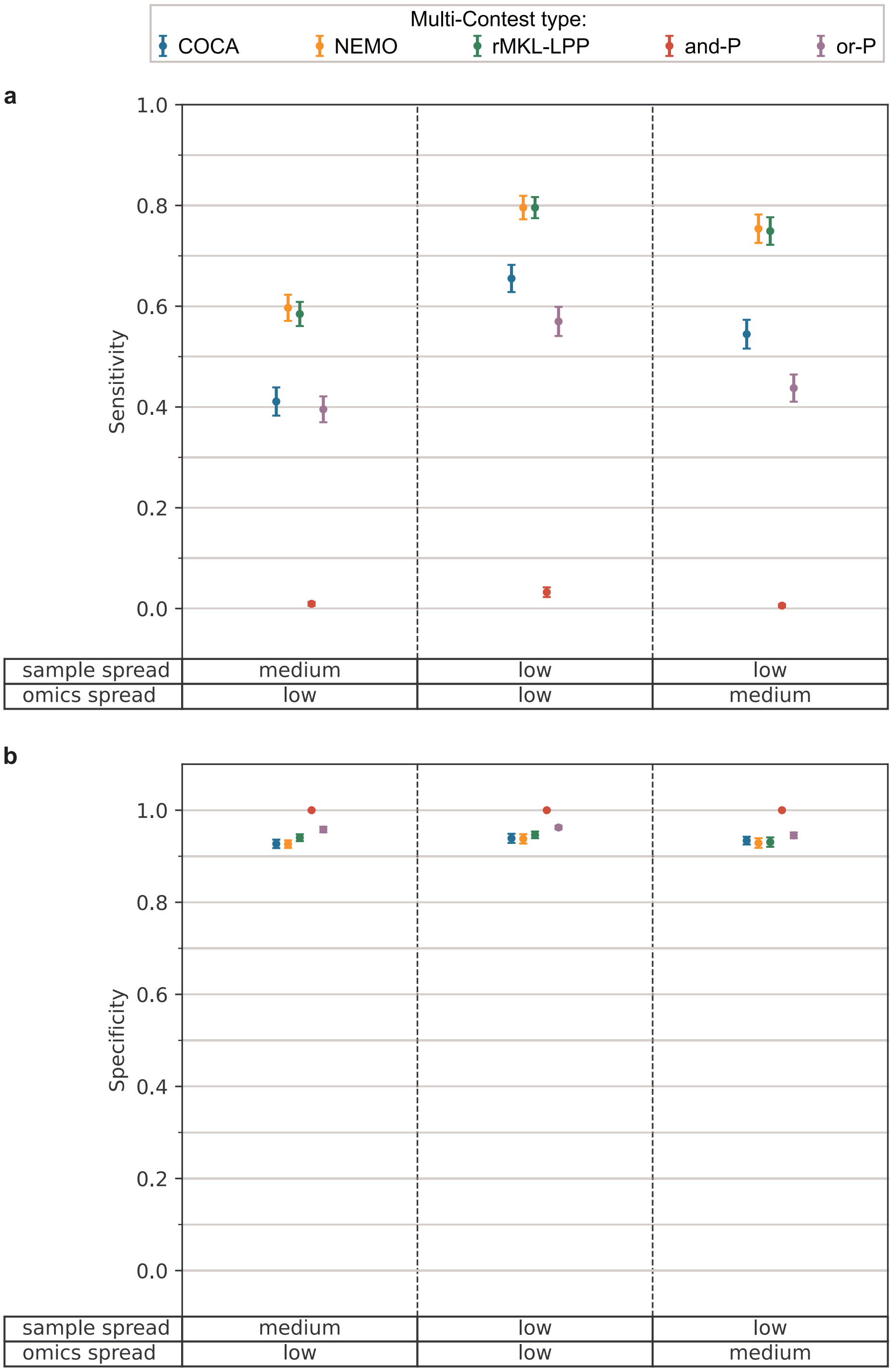


**Supplementary Figure 2:** Performance measures of Multi-Contest in simulated data sets with three cluster centers. The sensitivity (**a**) and specificity (**b**) are shown as means ± SEM from different strength of simulated confounding with ten omics (triplicates for each strength, total n = 24). Different combinations of sample spread and omics spread were used to test how the methods perform on less a less clear cluster structure.

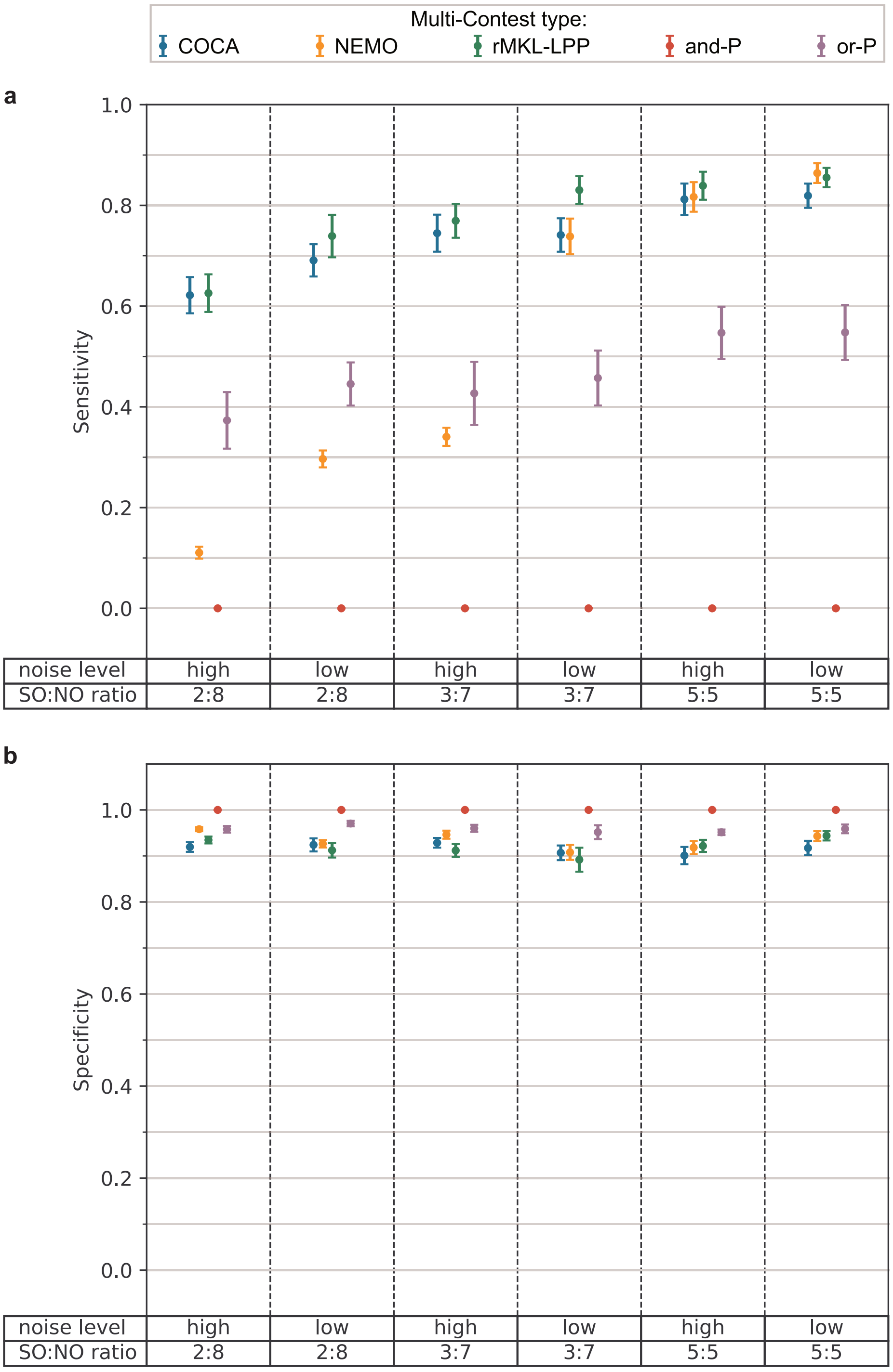


**Supplementary Figure 3:** Performance measures of Multi-Contest in simulated data sets containing signal omics and noise omics. The sensitivity (**a**) and specificity (**b**) are shown as means ± SEM from different strength of simulated confounding with ten omics (triplicates for each strength, total n = 12). The signal omics (SO) to noise omics (NO) ratio (SO:NO ratio) as well as the noise level used for the simulation are indicated at the bottom of the figure.
